# A synthetic lethal screen identifies HDAC4 as a potential target in MELK overexpressing cancers

**DOI:** 10.1101/2021.07.16.452653

**Authors:** Lin Zhou, Siqi Zheng, Fernando R. Rosas Bringas, Bjorn Bakker, Judith E. Simon, Petra L. Bakker, Hinke G. Kazemier, Michael Schubert, Maurits Roorda, Marcel van Vugt, Michael Chang, Floris Foijer

## Abstract

Maternal embryonic leucine zipper kinase (MELK) is frequently overexpressed in cancer, but the role of MELK in cancer is still poorly understood. MELK was shown to have roles in many cancer-associated processes including tumor growth, chemotherapy resistance, and tumor recurrence. To determine whether the frequent overexpression of MELK can be exploited in therapy, we performed a high-throughput screen using a library of *Saccharomyces cerevisiae* mutants to identify genes whose functions become essential when MELK is overexpressed. We identified two such genes: *LAG2* and *HDA3. LAG2* encodes an inhibitor of the SCF ubiquitin-ligase complex, while *HDA3* encodes a subunit of the *HDA1* histone deacetylase complex. We find that one of these synthetic lethal interactions is conserved in mammalian cells, as inhibition of a human homolog of *HDA3* (HDAC4) is synthetically toxic in MELK overexpression cells. Altogether, our work might provide a new angle of how to exploit MELK overexpression in cancers and might thus lead to novel intervention strategies.

## Introduction

Maternal embryonic leucine zipper kinase (MELK), a serine/threonine kinase, plays a dominant role in cell cycle regulation, proliferation, and apoptosis [1,2]. MELK is overexpressed in multiple human cancers, such as colorectal cancer [3], melanoma [4], basal-like breast cancer cells [5], and is a cell cycle-regulated gene regulated by E2F transcription factors [6]. Previous work has shown that MELK is involved in cytokinesis in Xenopus embryos and that MELK inactivation leads to cell division defects in this model [7–9]. Furthermore, RNA interference and inhibitor studies have shown that inhibiting MELK activity in cultured mammalian cells suppresses tumor cell growth [10–12]. In mouse models, overexpression of wild-type MELK protein leads to oncogenic transformation, which relies on its kinase activity. Conversely, *in vivo* inhibition of MELK activity through the inhibitor OTSSP167 in breast cancer xenograft models revealed that inhibition of MELK activity reduced the growth of basal-like cell breast cancer xenografts, but not of luminal cell breast cancer xenografts [11,13]. Similarly, OTSSP167 significantly improved the survival of mice in which A20 lymphoma cells were transplanted, which suggests that blocking MELK activity *in vivo* can also inhibit lymphoma progression [14]. Germline inactivation of MELK in mice does not yield an obvious phenotype during embryo development or in adult mice, indicating that MELK is dispensable for normal development [5]. Because of these combined observations, MELK is considered a highly selective cancer target.

Although previous studies provided evidence that MELK plays an important role in tumorigenesis, the precise role of MELK in tumor development has been challenged by more recent studies [13,15–17]. In particular, a study by the Sheltzer lab showed that CRISPR-mediated inactivation of MELK does not affect the fitness of triple-negative breast cancer cells, suggesting that OTSSP167 impairs cell division via off-target effects in these cells [15]. Another study showed no difference between the *in vivo* growth of xenograft MDA-MB-231 cells in which MELK was inactivated or not [18]. Therefore, the role of MELK in cancer is still not fully understood.

However, as MELK is so frequently overexpressed in human cancer [11,19–23], finding synthetic lethal interactions with MELK overexpression would still provide a powerful means to treat cancers that overexpress MELK. We therefore performed a high-throughput screen in *Saccharomyces cerevisiae* to identify genes that are required for yeast cell viability upon MELK overexpression. We followed up these findings in mammalian cells for two candidate genes, *LAG2* (encoding an inhibitor of the SCF ubiquitin-ligase complex) and *HDA3* (encoding a subunit of the *HDA1* histone deacetylase complex) [24,25], and found that inhibiting the latter renders mammalian cells sensitive to MELK overexpression. While MELK overexpression did not alter cell proliferation, concomitant inhibition of HDAC4 significantly attenuated the proliferation of MELK overexpressing mammalian cells. Altogether, our work shows that MELK overexpressing cells are sensitive to the inhibition of HDAC4, which may provide novel intervention strategies to target tumor cells that overexpress MELK.

## Materials and Methods

### Plasmids

To construct a doxycycline-inducible expression system for *MELK* overexpression, full-length cDNA encoding MELK was inserted between BamHI and NotI sites of three types of retroviral plasmids including the pRetroX-Tight-puro (Clontech), pRetroX-Tight-GFP-puro (containing N-terminal GFP) and pRetroX-Tight-BlastR (Puromycin exchanged by Blasticidin). The doxycycline-inducible pTRIPZ lentiviral shRNA vector targeting MELK (Catalog#: RHS4696-200691582) was purchased from Horizon Discovery. To generate inducible shRNA vectors targeting CAND1, oligonucleotides selected from the Sigma RNAi Consortium shRNA Library, annealed and directly ligated into gel-purified a EZ-tet-pLKO-BlastR [26] backbone digested with NheI and EcoRI. To generate a constitutive shRNA vector targeting *HDAC4*, oligonucleotides selected from the Sigma RNAi Consortium shRNA Library, annealed, and directly ligated into a gel-purified PLKO.1-puro (SHC001 Sigma-Aldrich) backbone digested with AgeI and EcoRI. All vectors were verified by Sanger sequencing. All primers are listed in Table 1.

**Table1.**
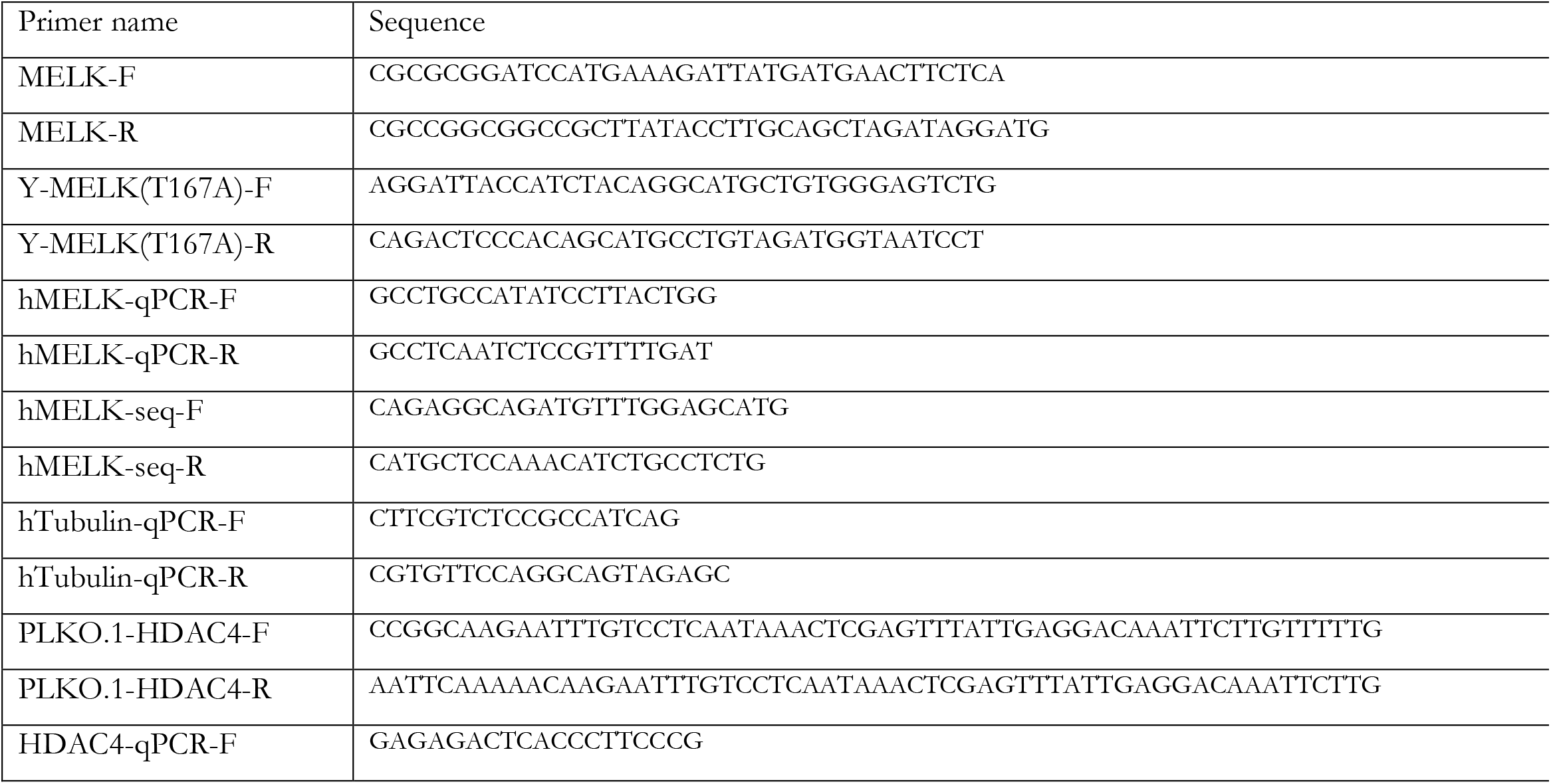
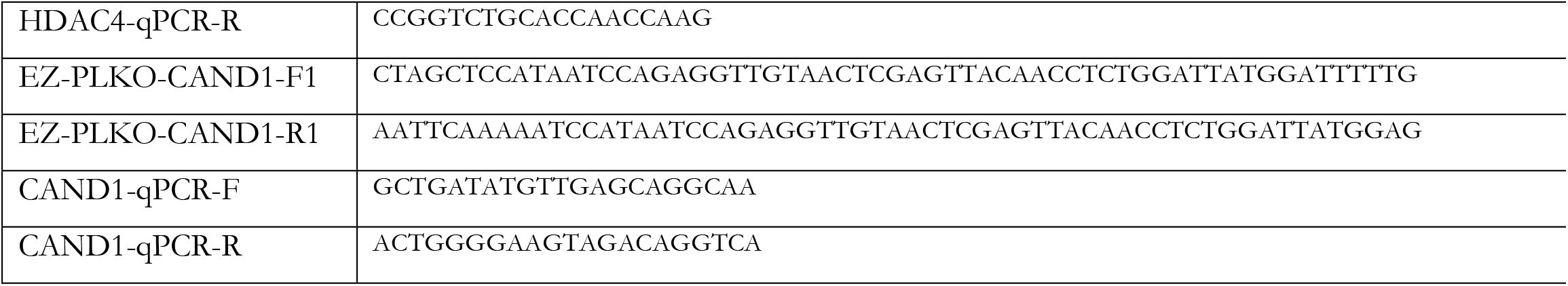
Primers used in this study.

To express human MELK in yeast, cDNA encoding human MELK was amplified by PCR using 5′-cgcgcggatccatgaaagattatgatgaacttctca-3′ forward and 5′-cgccggcggccgcttataccttgcagctagataggatg-3′ reverse primers. The resulting PCR product was cloned into a BamHI- and NotI-digested pSH380 plasmid. The kinase-dead MELK^T167A^ mutant was constructed using a Q5 site-directed mutagenesis kit (New England Biolabs) with 5′-aggattaccatctacaggcatgctgtgggagtctg-3′ forward and 5′-cagactcccacagcatgcctgtagatggtaatcct-3′ reverse primers.

### Cell lines and culture medium

Retinal pigmented epithelium (RPE)1, human mammary epithelial MCF10A cells, human mammary breast BT-20, MDA-MB-231 were purchased from the American Type Culture Collection (ATCC, Wesel, Germany). RPE1 p53 knockout cells were described earlier [27]. RPE1, BT-20 and MDA-MB-231 were cultured in DMEM (Gibco, Carlsbad, USA) with 10% fetal bovine serum (Invitrogen, Carlsbad, USA) and 50 U/µl Penicillin/Streptomycin solution. MCF10A cells were cultured in DMEM/F-12 (Gibco, Carlsbad, USA) supplemented with 5% horse serum (Invitrogen, Carlsbad, USA), 50 U/µl Penicillin Streptomycin, 100 ng/ml cholera toxin (Sigma), 20 ng/ml epidermal growth factor (Peprotech, Rocky Hill, CT, USA), 10 µg/ml insulin (Sigma), 500 ng/ml hydrocortisone (Sigma).

### Time-lapse Imaging

Time-lapse imaging was performed on a DeltaVision microscope (Applied Precision Ltd./GE). A total of 50,000 cells expressing pLNCX2 H2B-GFP were pre-seeded in 4-well imaging chambers (LabTech). Images were captured every 4 minutes with a 40X objective lens. Mitotic abnormalities from the overnight video were manually analyzed using softWoRxExplorer (Applied Precision Ltd./GE).

### Metaphase spreads

For metaphase spreads, cells were cultured up to 70% confluency with 10 µg/ml colcemid (Roche) for 3.5 h. Then cells were harvested and incubated in 75 mM KCl for 10 min at 37°C. Next, cells were fixed in MAA (methanol: acetic acid; 3:1) and dropped on glass slides, then stained with Vectashield-DAPI (Brunschwig Chemie bv.). Metaphase images were inspected on an Olympus microscope using a 60X lens.

### IncuCyte growth curves

Cells were seeded at a density of 40,000 cells/per well in 12-well plates. Cell growth was monitored every 2 h by the IncuCyte Zoom live-cell analysis system (Essen BioScience Ltd.). Cell density was quantified by IncuCyte Zoom 2018A software (Essen BioScience Ltd.).

### qRT-PCR analysis

Total RNA was isolated and purified from cultured cells with the RNeasy Mini kit (Qiagen). 1.5 µg of total RNA was reverse-transcribed to cDNA in a 20 µl mixture of random primers, 10x RT buffer, RNase inhibitor, and reverse transcriptase (New England Biolabs). cDNA was quantified by iTaq Universal SYBR Green supermix (Bio-Rad) on the LightCycler® 480 Instrument (Roche). Primers are listed in Table 1.

### Western Blot

Cells were harvested by trypsinization and lysed in elution buffer (150 mM NaCl, 0.1% NP-40, 5 mM EDTA, 50 mM HEPES pH7.5) containing complete protease inhibitor (Roche) for 30 minutes. Then the samples were centrifuged at 300g, at 4°C for 10 min to remove insoluble residues. 20 µg of each sample was loaded on 10% polyacrylamide gels. Then, proteins were transferred to polyvinylidene difluoride (PVDF) membranes. After blocking in Odyssey blocking buffer (Li-cor Biosciences) at 4°C for 30 min, the membrane was incubated overnight at 4°C with a primary antibody. Following incubation, the membrane was washed with 1X PBS containing 0,1% Tween 20 (Sigma) three times and incubated in secondary antibody for 1 hour at room temperature. The blots were detected by the Odyssey imaging system (Li-cor Biosciences). The protein bands were quantified with Image studio lite software (Li-cor Biosciences). We used the following antibodies: Anti-MELK antibody (ab108529, Abcam), α-Tubulin (ab7291, Abcam), anti-Mouse IRDye 680RD (P/N: 926-68072, Li-cor Biosciences), anti-Rabbit IRDye 800RD (P/N: 926-32211, Li-cor Biosciences).

### Colony-formation assay

To assess proliferation potential, cells were seeded at a density of 40,000 cells/per well in 12-well plates. After 5 days, cells were fixed with 4% formaldehyde and stained with crystal violet. For quantification of the staining, 1 ml of 10% acetic acid was added per well to elute crystal violet, and absorbance was measured at 590 nm wavelength with a synergy H1 plate reader.

### Yeast strains and methods

Standard yeast media and growth conditions were used [28,29]. For the synthetic dosage lethality screen, the pSH380, pSH380-MELK, and pSH380-MELK-kd plasmids were transformed into the yeast knockout (YKO) library [30] and a library of strains containing temperature-sensitive (ts) mutations of essential genes [31] using a high-throughput method called selective ploidy ablation (SPA) [32]. Replica-pinning steps were performed with a ROTOR HDA robot (Singer Instruments, Somerset, UK). Colony growth data were processed using the ScreenMill software suite [33] to identify colonies with reduced growth upon expression of MELK or MELK-kd. The screen was performed in biological triplicate, and in each screen, every strain was present in quadruplicate. A total of 255 putative “hits” were identified as having a growth defect upon expression of MELK at least once in the three replicate screens. A subset of these hits was validated by the transformation of the three plasmids into strains from the YKO or temperature-sensitive (ts) libraries using the LiAc-based method [34], and their growth was tested by spot assays on YP medium containing either glucose or galactose.

### Preparation of denatured extracts of yeast

10 ml of yeast culture grown to mid-logarithmic phase was centrifuged at 3000 rpm, 4 °C for 5 min. The cell pellet was resuspended in 1 ml of 20% TCA (Sigma) and transferred to a FastPrep tube, cells were pelleted again and resuspended in 200uL 20% TCA and glass beads for FastPrep were added. Yeast cells were lysed using FastPrep (MP Biomedicals) program: 6.0m/sec 40sec 1 cycle. 400uL of 5% TCA was added to the extract and the extract was collected by centrifugation at 1000 rpm, 4 °C for 1 min. Then the extract was centrifuged at 3000rpm for 10 minutes, and the pellet was resuspended in 100uL Laemmli buffer (150mM Tris pH6.8, 6% SDS, 30% glycerol, bromophenol blue dye, 7,7% B-ME) and 50uL 1M Tris Base and boiled for 5 min. To remove debris, the tube was centrifuged at 300rpm at 4 °C for 10 min, and the supernatant was transferred to fresh protein LoBind tubes. 20 µl of the denatured extracts were loaded on polyacrylamide gels for subsequent analysis by western blot.

### Aneuploidy and MELK expression correlation

We downloaded both gene expression and copy number segments from the TCGA breast cancer (BRCA) cohort via TCGAbiolinks tgca [35] and show the expression of MELK vs. the aneuploidy score. We computed the expression level of MELK using the variance stabilizing transformation of DESeq2 [36], and the aneuploidy score as the average deviation of the mean DNA copy number measurements along the genome, excluding X and Y chromosomes.

### DepMap analysis

For a large set of cancer cell lines, gene-level essentiality scores (CRISPR or RNAi based), copy number and mRNA expression data were obtained from DepMap release 21Q1, using the Broad Institute’s DepMap portal. Cell lines with fibroblast, teratoma, unknown, engineered or of non-cancerous origin were excluded from the analysis. Pearson correlation coefficients were computed using R 4.0.0 to identify associations between MELK copy number or mRNA expression and gene essentiality scores. To control false detection rate, p-values were corrected using Benjamini-Hochberg correction.

### Data availability Statement

Strains, cell lines and plasmids available upon request. The authors affirm that all data necessary for confirming the conclusions of the article are present within the article, figures, and tables.

## Results

### Identification of mutations sensitive to MELK overexpression

To identify genes that are required for cellular viability when MELK is overexpressed, we performed a synthetic dosage lethality screen in yeast. We chose to perform this screen in budding yeast, *Saccharomyces cerevisiae*, because of the speed with which one can perform genome-wide screens and because many aspects of eukaryotic cell biology, including mitosis and chromosome segregation, are highly conserved in this model organism. We first tested whether expression of human MELK or a kinase-dead variant of MELK (MELK-kd) is tolerated in yeast by introducing the human cDNAs under the control of the galactose-inducible *GAL1* promotor (Fig. 1A). We found that both human proteins are well tolerated and that their expression does not noticeably affect cell growth in yeast (Fig. 1B). We next introduced both plasmids into the yeast knockout library, consisting of ∼4800 yeast strains in which each nonessential gene is deleted [30], as well as a library of yeast strains containing temperature-sensitive alleles of essential genes [31], and determined potential synthetic lethal interactions by quantifying the colony size of each mutant strain with or without MELK or MELK-kd expression. We performed this screen three times and found 255 potential synthetic lethal interactions, *i*.*e*. mutant strains that grew slower when MELK was expressed, of which 214 grew slower only upon MELK, and not MELK-kd, expression (Supplemental Table S1). When we analyzed the gene ontologies of these cumulative candidate genes, we found the enriched GO terms to be mainly involved in the biological processes ‘cell cycle’, ‘mitotic nuclear division’ and ‘cell division’ (Supplemental Table S2), which very much resembles findings of a previous yeast screen studying processes involved in CIN [37].

**Figure 1.**
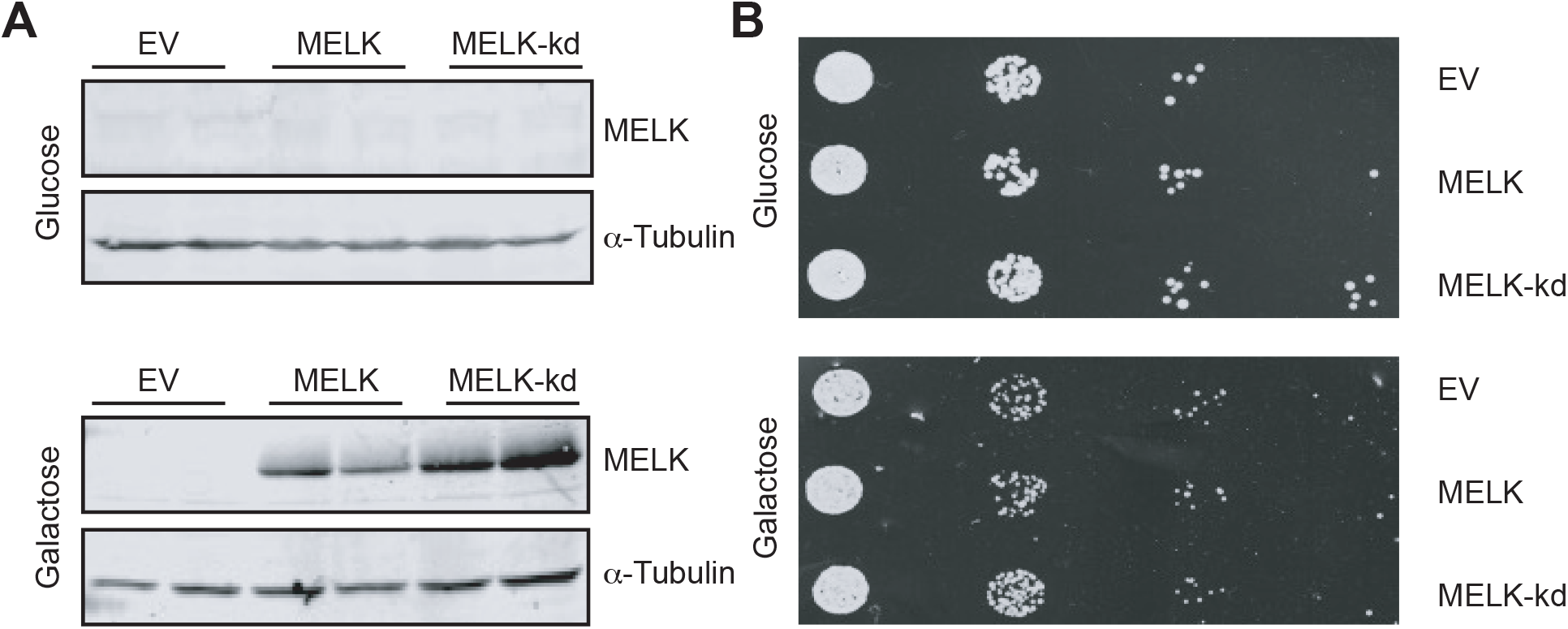
(**A**) MELK and MELK-kd were expressed from a galactose-inducible promoter in a wildtype yeast strain. (**B**) Serial tenfold dilutions of the indicated BY4741 yeast strains harboring either an empty vector, a plasmid expressing MELK, or a plasmid expressing MELK-kd were spotted onto YP medium-containing plates supplemented with either glucose or galactose.

### Altered expression of MELK does not impair mitotic fidelity in mammalian cells

Since our synthetic lethal screen in yeast suggested a role of MELK in the control of genomic integrity (Supplemental Tables S1, S2) in line with earlier observations in Xenopus laevis embryos [9], we next wanted to test whether MELK plays a role in maintaining genomic integrity in mammalian cells. We therefore determined whether MELK overexpression is correlated with aneuploidy in The Cancer Genome Atlas (TGCA) database [38]. Indeed, we found a strong positive correlation between MELK expression and aneuploidy (Fig. 2A), suggesting that MELK is somehow involved in genomic instability. To better understand how increased expression of MELK can lead to aneuploidy, we engineered a retina pigment epithelial (RPE1) cell line in which GFP-tagged MELK can be overexpressed under a doxycycline-inducible promotor. We first tested whether doxycycline addition indeed induced MELK-GFP expression and found that increasing doxycycline concentrations yielded higher MELK-GFP expression, which plateaued at 100 ng/ml of doxycycline. Quantification of the protein levels revealed that MELK-GFP was overexpressed 1.6-fold compared to endogenous MELK at this doxycycline concentration (Fig. 2B).

**Figure 2.**
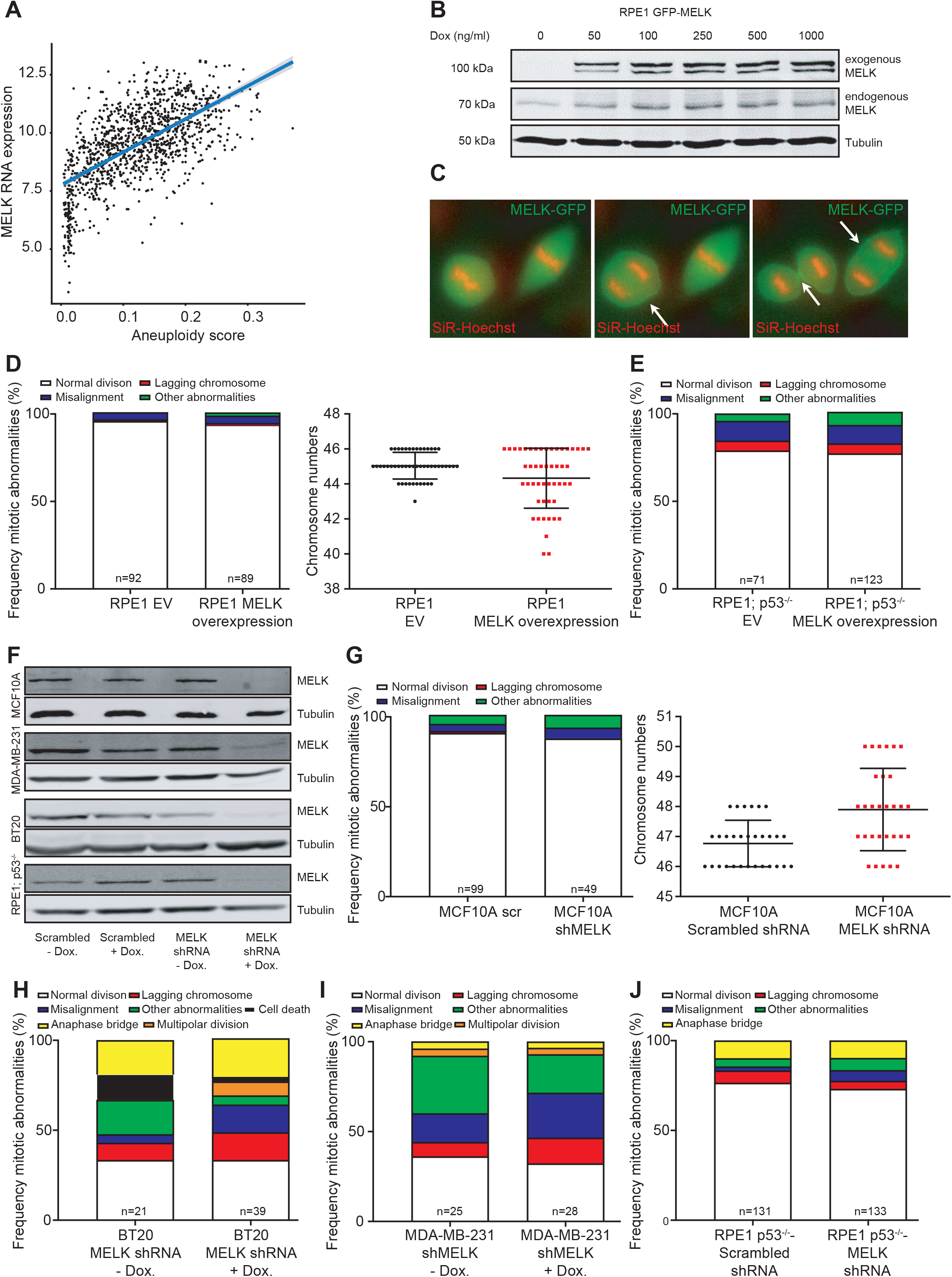
(**A**) Analysis of correlation between MELK expression and aneuploidy. (**B**) Western blot of doxycycline-induced GFP-tagged MELK and endogenous MELK in RPE1 cells. Tubulin serves as a loading control. (**C**) Time-lapse imaging still of mitotic RPE1 cells overexpressing GFP-tagged MELK showing that MELK localizes to the cleavage furrow during cytokinesis. Arrows indicate beginning (middle pane) and late cleavage furrow (right panel). (**D**) Frequency of mitotic abnormalities determined by time-lapse imaging (left panel) and quantification of karyotypes by metaphase spreads (right panel) in RPE1 cells with induced MELK overexpression. (**E**) Frequency of mitotic abnormalities determined by time-lapse imaging observed in MELK overexpressing RPE1 p53 knockout cells. (**F**) Western blots for MELK protein showing doxycycline-induced MELK knockdown in MCF10A, BT-20, MDA-MB-231 and RPE1 p53^-/-^cells compared to scrambled shRNA controls. Tubulin serves as a loading control. (**G**) Frequency of mitotic abnormalities determined by time-lapse imaging (left panel) and quantification of karyotypes by metaphase spreads (right panel) in MCF10A cells with MELK knockdown. (**H-J**) Frequency of mitotic abnormalities determined by time-lapse imaging observed in BT-20 cells (**H**), MDA-MB-231 cells (**I**) or RPE1 p53^-/-^cells with MELK knockdown (**J**).

As MELK was previously shown to be involved in cytokinesis in Xenopus embryos [20] and mammalian cells [21], we first monitored MELK-GFP localization throughout the cell cycle by time-lapse imaging. We found, in agreement with other studies [39–41], that MELK localizes to the mitotic cleavage furrow during cytokinesis in RPE1 cells (Fig. 2C). We therefore next introduced the live-cell nuclear stain SiR-DNA (Spirochrome) into doxycycline-inducible MELK-GFP RPE1 cells to label the chromatin and used time-lapse imaging microscopy to investigate the effect of MELK-GFP overexpression in mitosis. We found that overexpression of MELK did not impair mitotic fidelity (Fig. 2D), not even when we overexpressed MELK in p53^-/-^RPE1 cells (Fig. 2E). [27]. Conversely, we also tested whether the inactivation of MELK would alter chromosome missegregation rates. For this purpose, we engineered doxycycline-inducible shRNA vectors targeting MELK, which we transduced into MCF10A, BT-20, and MDA-MB-231 cells as the latter two breast cancer cell lines overexpress MELK with MCF10A cells as a control. Western blots confirmed that MELK levels were reduced in all three cell lines (Fig. 2F) compared to control cells expressing scrambled control shRNAs. We then introduced H2B-GFP by retroviral transduction in all shRNA-expressing cells and monitored cells by time-lapse imaging. This revealed that chromosome missegregation rates were altered in none of the MELK knockdown cell lines (Fig. 2G-J). Altogether, our results suggest that neither overexpression of MELK nor its depletion significantly alter mitotic fidelity, even in a p53-deficient background.

### MELK overexpression sensitizes human cells to HDAC4 knockdown

Since we did not find an effect of MELK overexpression or inhibition on chromosome segregation fidelity, we next decided to, instead of pursuing the role of MELK in chromosomal instability, pursue individual candidate genes that we had identified in the MELK synthetic dosage lethality yeast screen. We validated putative hits from our high-throughput screen by reintroducing the MELK expression plasmid, or the vector control, into a subset of the 255 putative hits, and spotting tenfold serial dilutions onto media containing glucose (non-inducing) or galactose (MELK-inducing). In total, only two of the 44 tested showed a robust growth defect upon MELK expression; *lag2Δ* is sensitive to MELK, but not MELK-kd, expression, while *hda3Δ* is sensitive to both (Fig. 3A). At present, we have no explanation for the high rate of false positives among our putative hits, especially given the enrichment of mitosis-related GO terms. One possibility is that MELK expression may have affected how the screen was performed, which involved the use of a plasmid-containing donor strain with 16 de-stabilizable centromeres and counterselectable chromosomes. Nevertheless, we decided to pursue Lag2 and Hda3 further as both have known human homologs, CAND1 and HDAC4, respectively. Lag2 is a longevity-assurance gene and its deletion significantly reduces the life span of yeast [42]. Lag2 was found to be a negative regulator of the SCF complex [43], a role that appears to be conserved in mammalian cells through its homolog CAND1 [44]. Hda3 is essential for the activity of the yeast histone deacetylase *HDA1* [45], similar to its mammalian homolog HDAC4, a Class II deacetylase that has a demonstrated role as a transcriptional repressor [46].

**Figure 3.**
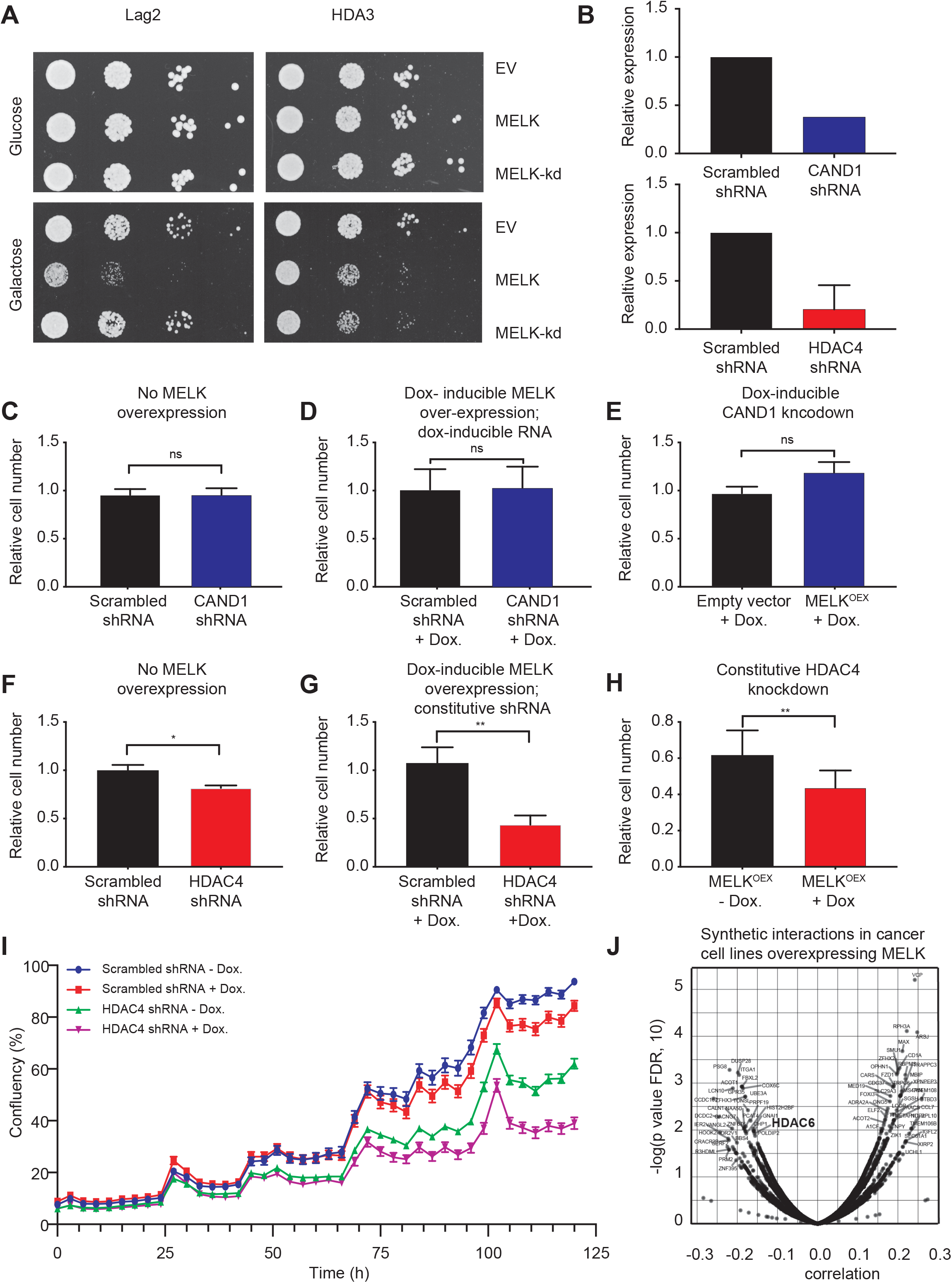
(**A**) Serial tenfold dilutions of indicated yeast strains harboring either an empty vector, a plasmid expressing MELK, or a plasmid expressing MELK-kd spotted onto YP medium plates containing either glucose or galactose. (**B**) RT qPCR quantification of CAND1 (upper panel) and HDAC4 (lower panel) transcript levels before and after the knockdown. Transcript levels were normalized to scramble shRNA (**C**) Quantification of colony formation potential of RPE1 cells with CAND1 knockdown (blue bar) compared to RPE1 cells expressing a control scrambled shRNA (black bar). (**D**) Quantification of colony formation potential RPE1 cells that overexpress doxycycline induced MELK with a control scrambled shRNA (black bar) or CAND1 knockdown (blue bar). (**E**) Quantification of colony formation potential of CAND1-depleted RPE1 cells expressing a doxycycline-inducible control vector (black bar) or doxycycline-inducible MELK (blue bar). (**F**) Quantification of colony formation potential of RPE1 cells with (red bar) or without (black bar) HDAC4 knockdown. (**G**) Quantification of colony formation potential of RPE1 cells that overexpress MELK with a control-scrambled shRNA (black bar) or with HDAC4 knockdown (red bar). (**H**) Quantification of colony formation potential of HDAC4-depleted RPE1 cells with (red bar) or without (black bar) doxycycline-induced MELK overexpression. (**I**) IncuCyte-determined proliferation curves showing growth of doxycycline-inducible MELK RPE1 cells expressing scrambled control shRNA with (red line) or without (blue line) doxycycline and doxycycline-inducible MELK RPE1 cells expressing an shRNA targeting HDAC4 with (purple line) or without (green line) doxycycline. (**J**) MELK mRNA expression correlates with RNAi knockdown lethality of HDAC6 in a large set of cancer cell lines (pearson correlation coefficient = -0.1572, *p* = 0.01186)

To validate whether the genetic interactions observed in yeast are conserved in human cells, we engineered shRNAs targeting CAND1 and HDAC4, which we transduced into MELK-overexpressing RPE1 cells. RT-qPCR confirmed reduced expression levels of CAND1 (∼60% knockdown) and HDAC4 (∼80% knockdown) (Fig. 3B). However, knocking down CAND1 did not alter cell proliferation in MELK-overexpressing RPE1 cells, nor control cells, suggesting that the synthetic interaction between MELK overexpression and CAND1 inhibition is not conserved in mammalian cells (Fig. 3C-E).

When we next interfered with HDAC4 expression in control RPE1 cells, we found that this decreases colony formation by approximately 20% (Fig. 3F), but that this growth defect is much stronger in RPE1 cells that overexpress MELK (Fig. 3G, growth reduction of 57%). Importantly, a direct comparison between HDAC4 knockdown cells with and without MELK overexpression showed that MELK overexpression significantly delayed proliferation, both in a colony formation assay (Fig. 3H) as well as in an IncuCyte proliferation assay (Fig. 3I). Finally, we wanted to test whether our findings can be extrapolated to human cancer cell lines as well. We therefore downloaded expression data and copy number status of MELK from the Cancer Cell Line Encyclopedia (DepMap release 21Q1, also see Methods) and tested which genes are essential in cells overexpressing MELK from RNAi screens. We found that, among other genes, inactivation of HDAC6, another type II HDAC [47], becomes more toxic when MELK is highly expressed, evidenced by a negative correlation between gene essentiality and MELK expression (Fig. 3J), further strengthening our finding that inhibition of type II HDACs is particularly toxic in MELK overexpressing cells. Together, our results demonstrate that inhibition of HDAC4 activity impairs the proliferation of cells that overexpress MELK more than cells that do not overexpress MELK, which might reveal an interesting targetable vulnerability of cancers that overexpress MELK.

## Discussion

MELK is frequently overexpressed in cancer, which makes it a promising target in cancer therapy. However, as MELK is an E2F target gene [6], its increased expression could well be a side-effect of increased activity of E2F activity in cancer cells. This is well in line with the observation that MELK expression is positively correlated with the expression of several other proliferation markers such as MCM2, KI67, PCNA, CCNB1, and TOP2a [18]. Therefore, it is unclear whether and how increased expression of MELK contributes to the fitness of cancer cells. While we do find that human cancers that overexpress MELK are significantly more aneuploid, we failed to find any direct effects of MELK overexpression on mitotic fidelity or cell proliferation. Our findings are in agreement with previous findings showing that MELK inactivation in triple-negative breast cancer cells does not alter proliferation rates [15–18].

We found that deletion of *LAG2* and *HDA3* renders yeast cells sensitive to the expression of MELK. However, the toxic interaction between loss of the *LAG2* homolog CAND1 and MELK overexpression was not conserved in mammalian cells. A possible explanation for this is that *LAG2* functions are not fully conserved between *S. cerevisiae* and mammals. Thus, the toxic interaction observed in yeast may be caused by a non-conserved role of *LAG2*. Alternatively, human cells may have additional pathways that can buffer the loss of CAND1. Finally, our shRNA vector-only reduced CAND1 by 60%, which might have been sufficient to recapitulate the phenotype observed in yeast, in which the Lag2 gene was completely inactivated. Depletion of the *HDA3* homolog HDAC4 histone deacetylase, on the other hand, did decrease the proliferation of MELK overexpressing cells. Importantly, this growth-inhibiting effect of HDAC4 inhibition was much stronger in MELK overexpressing cells compared to control cells. HDAC4 is a transcriptional repressor and its inhibition was found to reduce the growth of colon cancer cells through upregulation of p21 [45,46,48]. While MELK indeed is typically highly expressed in colon carcinoma cell lines [49], further work is needed to determine whether this observed sensitivity to HDAC4 inhibition depends on MELK expression. Even though our work does not provide further mechanistic insight into the nature of the interaction between MELK and HDAC4, it is intriguing that MELK was identified as a *bona fide* interaction partner of HDAC4 in the BioGRID database [50] and that that HDAC6 inactivation becomes increasingly toxic with overexpression of MELK in cancer cell lines in the CCLE database (Fig. 3J). Since both HDAC6 and HDAC4 have been shown to facilitate DNA damage repair in glioblastoma [51], the synthetic interaction between HDAC4 and HDAC6 with MELK could indicate a role for MELK in DNA damage as well. While our work provides functional evidence for a synthetic interaction between MELK overexpression and type-II HDACs, further experiments are needed to map the molecular mechanism underlying these interactions. Altogether, our work could provide a new angle of how to target MELK-overexpressing cancers and might thus lead to novel intervention strategies in the future.

## Acknowledgments

We are grateful to the members of the Foijer and Chang lab for fruitful discussions. This work was funded by a Chinese Scholarship Council fellowship to Zhou, a UMCG Cancer Research Fund (KRF) grant to Zhou, and a Dutch Cancer Society grant (2015-RUG-7833) to Foijer.

